# Scalable recombinase-based gene expression cascades

**DOI:** 10.1101/2020.06.20.161430

**Authors:** Tackhoon Kim, Benjamin Weinberg, Wilson Wong, Timothy K. Lu

**Affiliations:** Research Lab of Electronics, Massachusetts Institute of Technology, Cambridge, Massachusetts, USA; Chemical Kinomics Research Center, Korea Institute of Science and Technology, 5 Hwarangro 14-gil, Seongbuk-gu, Seoul 02792, Republic of Korea; Department of Biomedical Engineering and Biological Design Center, Boston University, Boston, Massachusetts, USA

## Abstract

Temporal modulation of multiple genes underlies sophisticated biological phenomena. However, there are few scalable and generalizable gene circuit architectures for the programming of sequential genetic perturbations. We describe a modular recombinase-based gene circuit architecture, comprising tandem gene perturbation cassettes (GPCs), that enables the sequential expression of multiple genes in a defined temporal order by alternating treatment with just two orthogonal ligands. We used tandem GPCs to sequentially express single-guide RNAs to encode transcriptional cascades and trigger the sequential accumulation of mutations. We built an all-in-one gene circuit that sequentially edits genomic loci, synchronizes cells at a specific stage within a gene expression cascade, and deletes itself for safety. Tandem GPCs offer a multi-tiered cellular programming tool for modeling multi-stage genetic changes, such as tumorigenesis and cellular differentiation.

## Main Text

Complex cellular tasks, such as those executed during normal development and tumorigenesis, are coordinated by multiple gene regulatory events operating across various time-scales. For example, the differentiation of cells into specific subtypes involves highly orchestrated transcriptional programs whose temporal regulation is controlled by transcriptional cascades of multiple genes (*1*). Tumorigenesis involves the mutation of key tumor suppressors and proto-oncogenes, and different temporal orders of these mutations may change disease progression (*2, 3*). Only a few gene expression cascades have been designed to model sequential events in tumorigenesis (*4*). However, these cascades require multiple independently integrated transgenes and may take months or even years to establish *in vivo*.

One of the central difficulties in programming gene expression cascades is scalability. Traditionally, cascades composed of multiple inducible gene expression systems require many distinct regulatory proteins that respond to different ligands (e.g., TetR for tetracycline) (*5*). Thus, the number of constitutively expressed transgenes increases linearly with the number of independent genetic tasks in the gene expression cascade. Such cascade designs can be large and challenging to implement and can be detrimental to cell viability because of resource competition at the transcriptional and translational level (*6–8*). Furthermore, the number of well-validated and well-tolerated inducible systems available for use in mammalian cells and in animal models is limited.

We overcame these limitations by designing an array of recombinase-based modules with memory that only require two distinct inducers; these modules reduce the constitutive expression of the transgenes needed to encode cascades. Each module performs two tasks: (1) the expression of payload gene(s), and (2) the self-termination of payload gene expression, followed by the expression of the next payload gene(s), in response to an inducer (Fig. 1A). To implement these features, we designed a compact, recombinase-based gene circuit called the gene perturbation cassette (GPC) (Fig. 1B). The GPC is composed of three parts: (1) a split recombinase that is activated in response to chemically induced dimerization (CID) by gibberellin (GIB) or abscisic acid (ABA) (*9–11*), (2) payload gene(s) expressed in the same mRNA transcript as the split recombinase and terminator, and (3) recombinase recognition sites that flank all genetic elements of the GPC. By placing a constitutively active promoter upstream of the GPC, the split recombinase and payload gene are expressed until the cognate ligand activates the recombinase, which excises the entire cognate GPC. This terminates recombinase activity and payload gene(s) expression and leads to expression of the next downstream GPC. This design provides an advancement in the scalability of programming gene expression cascades because temporal gene expression can be induced by simply alternating the exposure of the cells to just two ligands (ligand #1 for odd-numbered stages, and ligand #2 for even-numbered stages). The length of the gene expression cascade is limited only by the number of orthogonal recombinases; so far, nearly a dozen orthogonal recombinases have been confirmed to be active in mammalian cells (*12*). Furthermore, this architecture further minimizes cellular burden because only the recombinase and payload gene(s) in the proximal GPC are expressed at any given time.

**Figure 1.**
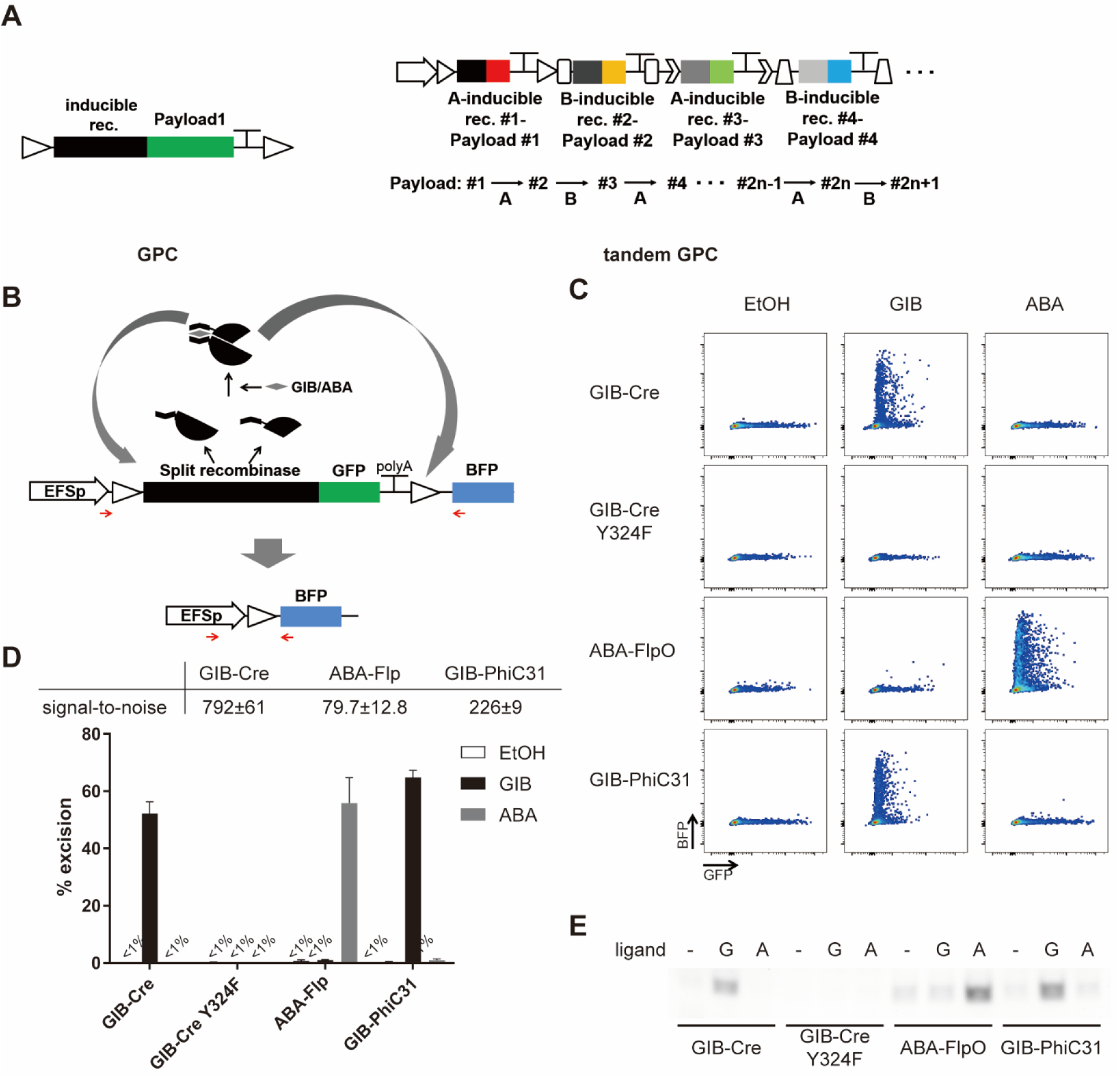
Design and validation of the Gene Perturbation Cassette (GPC). (A) Schematics of a single-unit GPC (left) and the tandem GPC circuit for programming gene expression cascades (right). (B) A single GPC enables the switching of the expressed gene from GFP to BFP. Red arrows indicate the primers used to detect the excision event in (E). (C) Representative flow cytometry plot for cells transfected with the single GPC architecture described in (B), implemented with the indicated split recombinase and treated with the indicated ligands for 24 hours. (D) Quantification of the results in (C) (n=3, mean ± s.d.). (E) PCR validation of excision upon ligand treatment using primers depicted as red arrows in (B). G: GIB, A: ABA.

Individual GPCs were optimized by implementing them in a simple two-stage cascade that switches expression from GFP (stage 1) to BFP (stage 2) upon ligand treatment and GPC excision (Fig. 1B). Split Cre, PhiC31, and Flp integrases were split at specific sites that give the highest signal-to-noise ratio, defined as the ratio of excision-induced BFP-positive cells in the presence and absence of CID ligand (Fig. S1A). We then appended an additional nuclear localization signal to these proteins for added recombinase activity (Fig. S1B). Furthermore, we selected the most efficient polyadenylation signals to encode at the end of each GPC in order to block leaky expression of downstream GPCs (Fig. S1C). Optimized GPCs, including GIB-activated Cre (GIB-Cre), GIB-activated PhiC31 integrase (GIB-PhiC), and ABA-activated Flp (ABA-Flp), had signal-to-noise ratios of 80~792 (Fig. 1C-D). We confirmed that CIDs by GIB and ABA are orthogonal to each other (Fig. 1C, D). Payload gene switching from GFP to BFP was due to recombinase activity, as a GPC with an inactive mutant Cre (Cre Y324F) (*13*) failed to switch the expressed gene to BFP (Fig. 1C, D). Excision-dependent payload gene switching was confirmed by detection of an excision-specific PCR product (Fig. 1E).

We next designed a gene expression cascade consisting of the tandem GPC array GIB-Cre → ABA-Flp → GIB-PhiC (Fig. 2A). Only the GPC directly proximal to the upstream promoter was expressed, consistent with our initial design in Fig. 1A. Thus, the initial GIB treatment, which activated GIB-Cre in the first GPC, did not excise the downstream GIB-PhiC GPC because the downstream GPCs were not initially expressed; this feature of the architecture allows GIB to be used again at stage 3 within the same cascade. We optimized the CID ligand treatment schedule to be 48 hours of each ligand with a 12-hour gap in between them (Text S1 and Fig. S2A). We used gibberellin A_4_ (GA_4_) (*14*), instead of the commonly used acetoxymethyl-gibberellin A_3_ to activate recombinase, as cells were readily permeable to GA_4_ and as it was rapidly cleared after removal from the growth medium (see Text S1 and Figs. S2B-C for further discussions).

**Figure 2.**
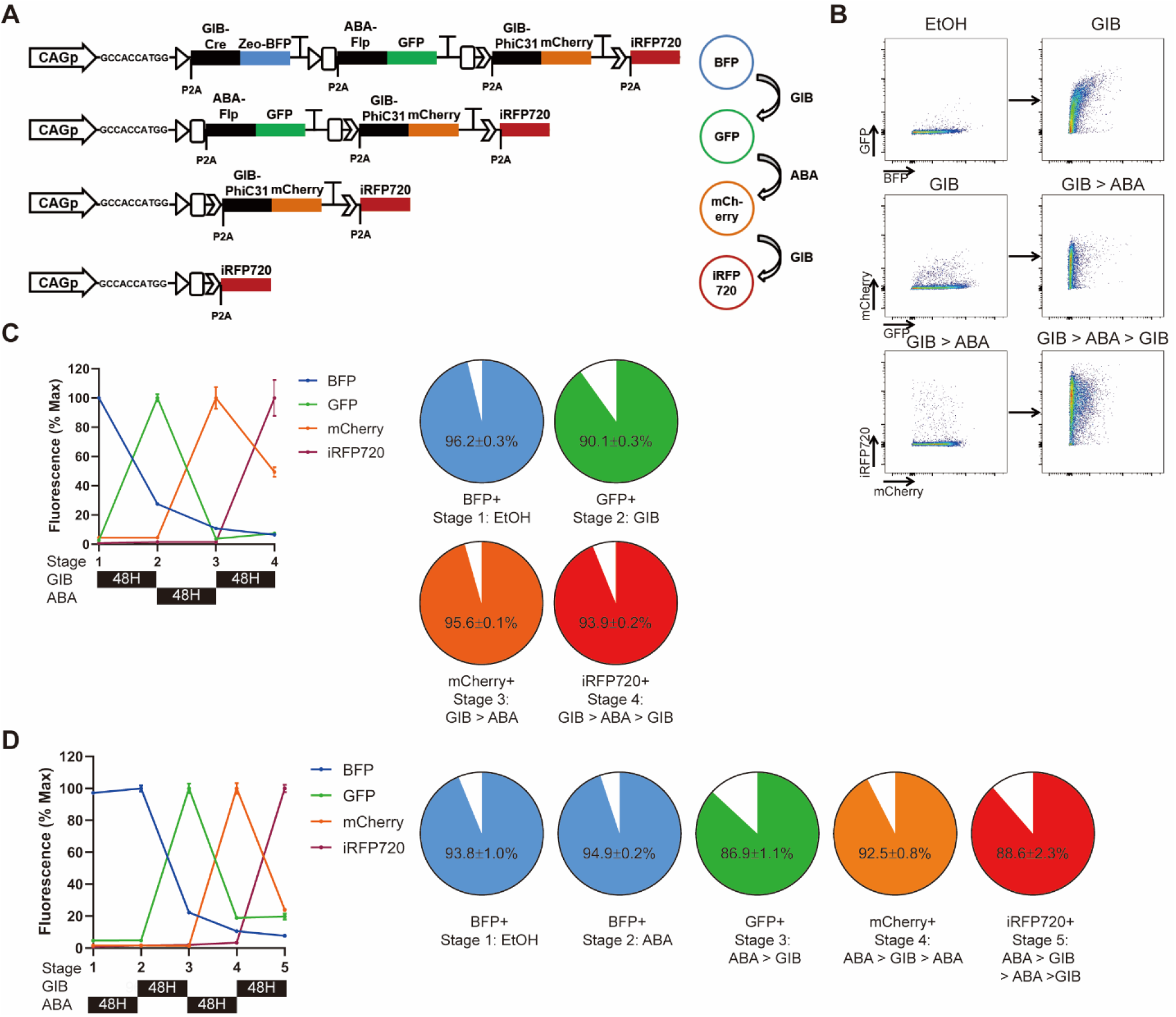
Design and validation of the tandem GPC gene circuit. (A) Tandem GPCv2-Fluor gene circuit, used in (B)-(D). Alternating treatments of GIB and ABA result in excision of GPCs and payload gene switching from BFP to GFP, mCherry, and finally iRFP720. Zeocin resistance gene is placed in the GIB-Cre GPC for positive selection of the gene circuit. (B) Representative flow cytometry plot of HEK 293T cells carrying tandem GPCv2-Fluor treated with the indicated sequence of CID ligands. (C) Quantification of mean fluorescence and average fraction of cells expressing the indicated fluorescent protein at each stage of the gene expression cascade (n=3, mean ± s.d.). (D) Quantification of mean fluorescence and average fraction of cells expressing the indicated fluorescent protein after each indicated sequence of ligand treatment (n=3, mean ± s.d.).

Excision by recombinases leaves a scar sequence consisting of recombinase-recognition sites (Fig. S2D). Therefore, in our original tandem GPC (version 1), where the the start codon was placed within each GPC, sequential excision of the tandem array of GPCs inevitably leads to lengthening of the scar upstream of the GPC to be expressed. Longer scars in the 5’ untranslated region (5’UTR) may form secondary structures that affect translation efficiency (*15*). To address this issue, we created a version 2 tandem GPC (Fig. S2E) with an initiation codon and a Kozak sequence between the promoter and the GPCs so that the scar sequence did not contribute to the 5’UTR but was actually translated. A self-cleaving 2A sequence was appended at the 5’ end of each split recombinase gene to reduce the possibility of the polypeptide from the scar sequence interfering with CID or recombinase activity. Consistent with our expectations, version 2 of the tandem GPC gene circuit had better excision efficiency than version 1, yielding a more robust gene expression cascade (Fig. S2E). All gene expression cascades described hereafter were based on version 2 of the tandem GPC design.

We next tested the tandem GPC gene circuit named tandem GPCv2-Fluor, which sequentially expresses four fluorescent proteins in HEK293T cells (Fig. 2A). Tandem GPCv2-Fluor consists of GIB-Cre-BFP, ABA-Flp-GFP, GIB-PhiC31-mCherry, and iRFP720 (*16*) so that alternating treatment with GA_4_, ABA, and GA_4_ induces the sequential expression of BFP, GFP, mCherry, and iRFP720. All payload genes were tagged with the PEST sequence (*17*) to increase their turnover rate so that fluorescence levels would more accurately reflect gene expression from each GPC. PiggyBac transposase was used to integrate large (>15kb) gene circuits into mammalian host genomes (*18*). Sequential treatment with GA_4_, ABA, and GA_4_, each for 48 hours, induced the gene expression cascade BFP→GFP→mCherry→iRFP720 in up to 96% of the cells (Fig. 2B-C).

We further assessed the robustness of the gene expression cascade by changing the ligand treatment schedule or the order of ligand treatment. We first tested whether cells with tandem GPCv2-Fluor maintained the memory of specific stages within the cascade. We did this by treating cells with GA_4_ to switch the payload gene to GFP and then maintaining them in the absence of CID ligand for 8 days. >80% of the cells retained the correct memory of the stage in the cascade; that is, they preserved GFP expression and the capability to switch the payload gene to mCherry in response to ABA treatment (Fig. S2F). We observed a minor (<10%) decrease in %GFP+ cells and premature mCherry expression in the absence of ABA. Point mutations that mitigate the tendency of split recombinase fragments to spontaneously reconstitute in the absence of CID ligand (*19*) will further reduce leakage and improve the fidelity of memory. We also treated cells carrying a tandem GPCv2-Fluor with CID ligands in a different temporal order: ABA→GA_4_→ABA→GA_4_. As expected, initial ABA treatment did not induce payload gene switching, nor did it disrupt the ability of tandem GPCv2-Fluor to execute the gene expression cascade upon subsequent GA_4_→ABA→GA_4_ treatment (Fig. 2D).

Beyond expressing genes, tandem GPCs can be used for the sequential expression of single-guide RNAs (sgRNAs) to leverage the versatility of CRISPR-Cas9 for driving transcriptional programs (tandem GPCv2-CRISPRa) or sequential genome editing (tandem GPCv2-CRISPR). Payload genes in GPCs are expressed together with the recombinase from an RNA polymerase II promoter; this contrasts with the standard RNA polymerase III promoters used to drive sgRNA expression in most studies. Thus, we compared several strategies (*20–23*) for RNA polymerase II-driven sgRNA expression by flanking an sgRNA with RNA cleavage sequences using an assay in which the sgRNA targets dCas9-VPR to an artificial sgRNA-responsive promoter to induce mCherry expression (Fig. 3A, left) (*24*). sgRNA flanked by 20nt core sequences for Csy4-mediated cleavage (“20nt core”), devoid of the 8nt “handle” sequence for Cas complex assembly (*25*), resulted in the strongest mCherry expression (Fig. 3A). Thus, in our tandem GPCv2-CRISPRa and tandem GPCv2-CRISPR circuits, the sgRNA was flanked by this 20nt core to induce the efficient liberation of sgRNA, while the remaining mRNA coding for the recombinase and lacking a poly-A tail was stabilized by appending a MALAT1 lncRNA triple helix (*26*).

**Figure 3.**
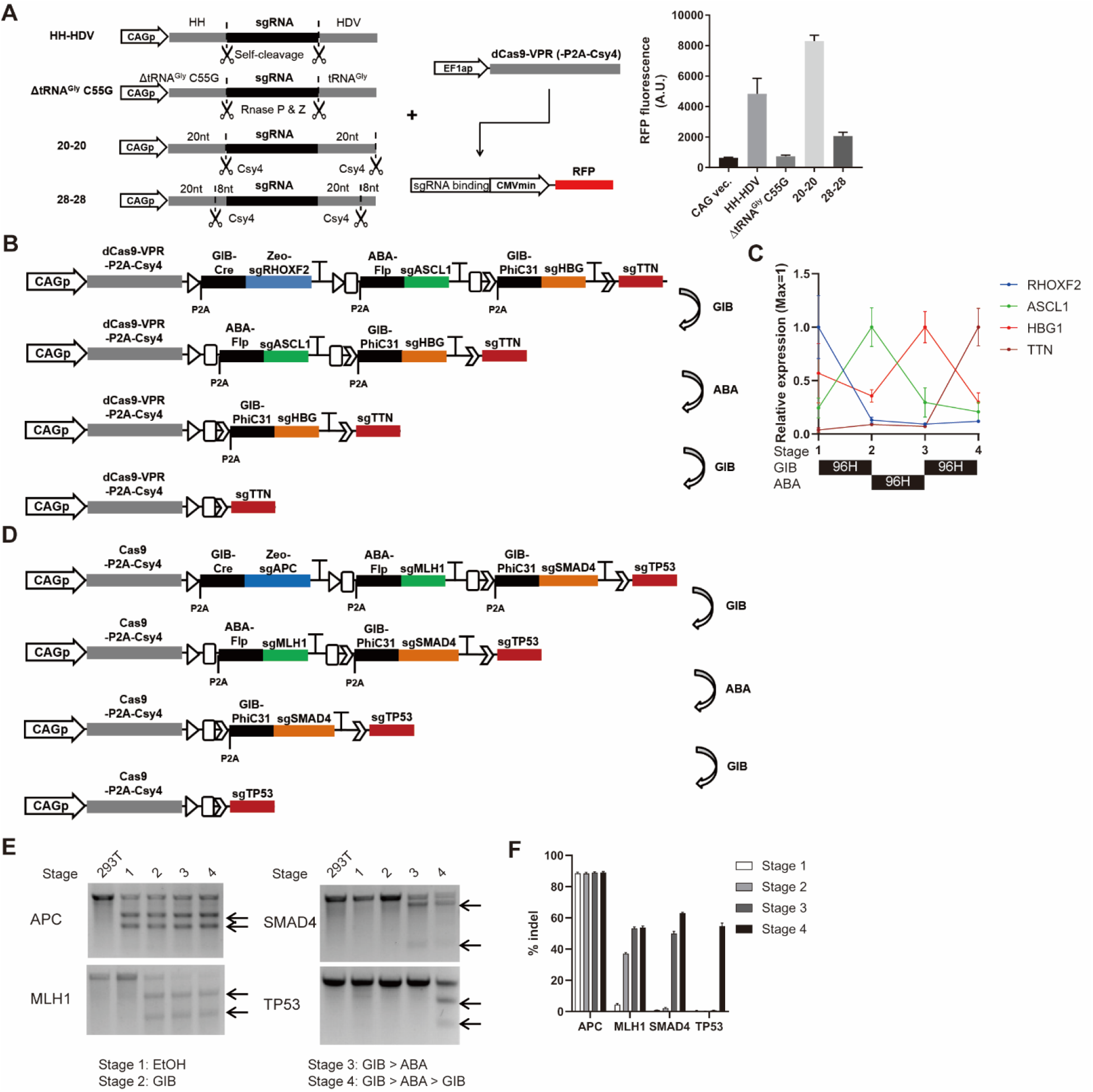
Sequential transcriptional cascade and gene editing by tandem-GPC-CRISPRa and tandem-GPC-CRISPR, respectively. (A) (Left) Schematics of different architectures for RNA Pol II-driven sgRNA expression and reporter construct for sgRNA activity. (Right) Activity of RFP sgRNA reporter with indicated RNA Pol II-driven sgRNA format (n=2, mean ± s.d.). (B) Tandem GPCv2-CRISPRa schematics. (C) Quantification of sgRNA target genes after each indicated sequence of ligand treatment (n=3, mean ± s.d.). (D) Tandem GPCv2-CRISPR schematics. (E) Validation of sequential gene mutations using T7 endonuclease assay. (F) Quantification of gene editing in (E) by next generation sequencing (NGS) (n=3, mean ± s.d.).

We next implemented the optimized tandem GPCv2-CRISPRa, which activates the transcriptional cascade of endogenous genes RHOXF2, ASCL1, HBG1, and TTN upon sequential treatment with GA_4_→ABA→GA_4_. We first examined the kinetics of decay of sgRNA after ligand treatment and excision of the GPC, and expression of the payload sgRNA in the next GPC. To this end, HEK293T cells carrying the tandem GPCv2-CRISPRa circuit were treated with GA_4_. Then we measured the kinetics of expression of RHOXF2 and ASCL1, which are the target genes of the sgRNAs expressed in the first and second GPC, respectively. In line with a previous report measuring sgRNA half-lives in cells (*27*), the expression levels of RHOXF2 and ASCL1 genes quickly changed, approaching equilibrium by 96 hours (Fig. S3). Therefore, we treated cells carrying tandem GPCv2-CRISPRa with CID ligands for 96 hours to transition between each stage in this cascade. As expected, tandem GPCv2-CRISPRa actuated a transcriptional cascade of RHOXF2, ASCL1, HBG1, and TTN in a temporally regulated manner (Fig. 3B, C).

We similarly implemented an sgRNA expression cascade for the sequential mutation of multiple genes by Cas9 to highlight the potential uses of this gene circuit for modeling the multi-stage nature of tumorigenesis (*28*). To this end, we made tandem GPCv2-CRISPR, a circuit that expresses a cascade of sgRNAs to kncokout key tumor suppressor genes in colorectal cancer. These genes, APC, MLH1, SMAD4, and TP53, are known to undergo sequential cancer-causing mutations (*3*). A T7 endonuclease assay and next-generation sequencing analysis revealed that sequential indel mutations were triggered in these four genes in 54~89% of the cells in response to the proper sequence of inducers (Fig. 3D-F).

The robustness of gene expression cascades is prone to decay with the number of stages within the cascade because recombinases are not 100% efficient. Furthermore, removal of the gene circuit after completion of the cascade may be desired for safety. To address these points, we created the AttP-tandem GPCv2-CRISPR gene circuit to encode a gene expression cascade of sgAPC→sgMLH1→Puro^R^-sgSMAD4, with AttP sites for PhiC31 integrase placed at the upstream end of the gene circuit (Fig. 4A). This all-in-one gene circuit sequentially generates indel mutations at APC, MLH1, and SMAD4 loci and then, by puromycin selection, synchronizes the cells that are at a specific stage (in this case, stage 3) by removing any cells that are at other stages (in this case, stages 1 or 2) in the gene expression cascade (Fig. 4B). After synchronization, GA_4_ treatment induces self-deletion of the AttP-tandem GPCv2-CRISPR gene circuit via recombination by PhiC31 integrase to prevent any further unwanted mutations. We confirmed the sequential accumulation of indel mutations in the sgRNA target loci (Fig. 4C, D). ~79% of the cells were synchronized at stage 3 in the gene expression cascade in response to puromycin selection (Fig. 4E). Synchronization removed cells that lacked indel mutations in the three loci due to inefficiencies in recombinase-mediated excision and subsequent gene expression cascade failure. The result of this synchronization was an increase in indel frequencies for APC, MLH1, and SMAD4 genes from 28~62% to 46~84%. Finally, the activation of GIB-PhiC removed the entire gene circuit in 67% of the cells (Fig. 4F), leaving cells that had specific indel mutations but were devoid of any exogenous genes, including Cas9, being expressed in the gene circuit.

**Figure 4.**
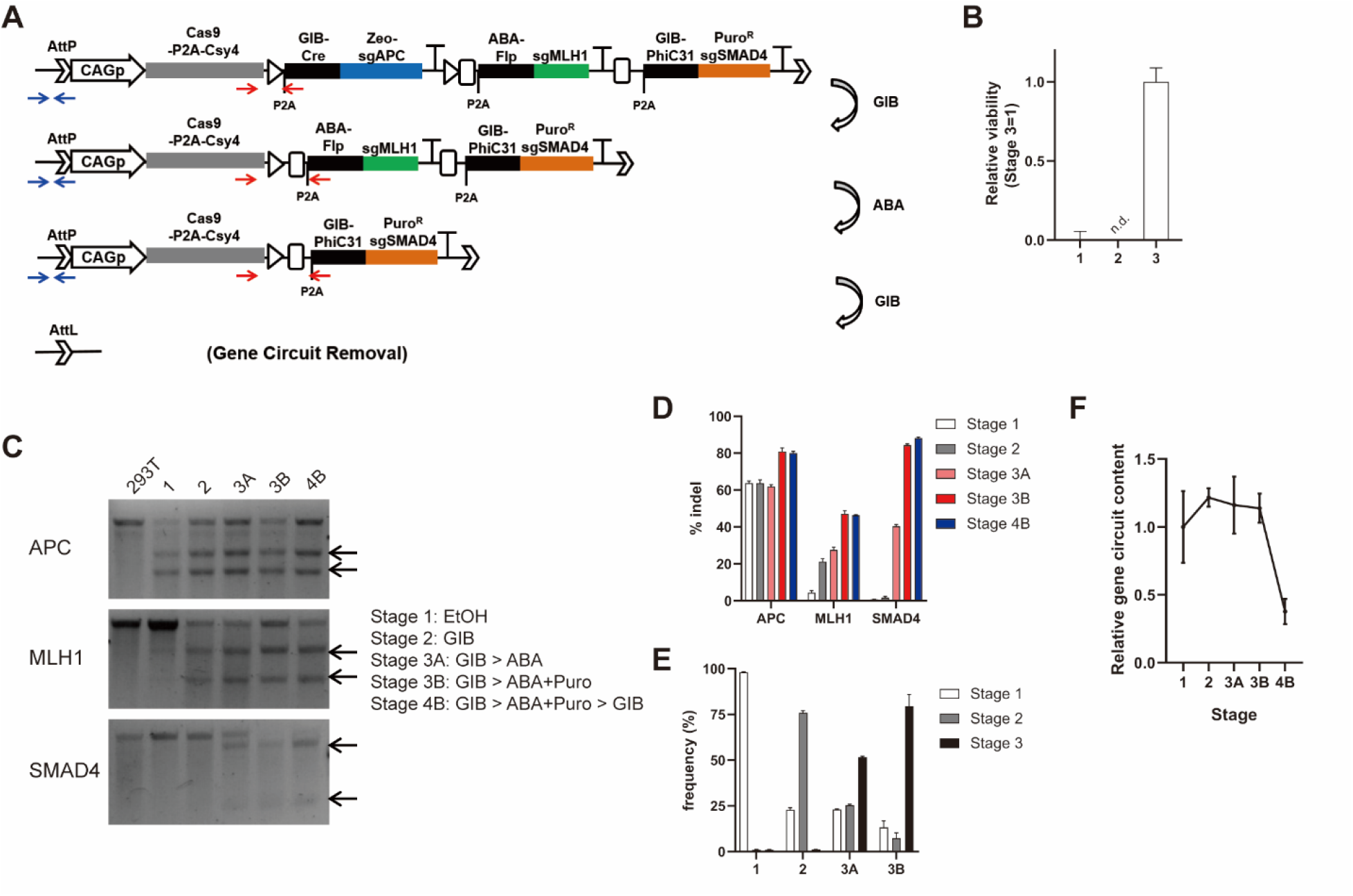
Sequential actuation, synchronization, and gene circuit removal by AttP-tandem GPCv2-CRISPR. (A) AttP-tandem GPCv2-CRISPR schematics. Red arrows indicate primers used for amplifying the scar sequence to identify the stage within the cascade, used in (E). Blue arrows indicate primers used for assessing gene circuit removal in (F). (B) Viability of cells under puromycin selection at specific stages in the cascade. (C) Validation of sequential gene mutations using T7 endonuclease assay. (D) Quantification of gene editing in (C) by NGS (n=3, mean ± s.d.). (E) NGS quantification of frequency of cells at each stage in cascade (n=3, mean ± s.d.). (F) qPCR validation of gene circuit removal (n=3, mean ± s.d.).

In summary, we demonstrated highly robust and scalable gene expression cascade circuits enabled by alternating treatment with two orthogonal ligands. We expect that gene circuits with diverse payloads in any arbitrary order will work in a “plug and play” manner, particularly if the flexibility of CRISPR-based transcriptional regulation and knockouts is leveraged. This feature thus enables the construction of gene expression cascades without intensive optimization.

We anticipate that this gene circuit will be a useful tool for research and medicine. This circuit can be used to model cancer, both *in vitro* and *in vivo*, because it can be designed to closely mimic not only the multitude of mutations in human cancers but also the temporal order in which they occur. Similarly, we envision that this gene circuit can be applied for the efficient direct reprogramming of one cell type to another because it can be designed to reproduce the transcriptional cascades that occur in natural differentiation processes. This application should be useful in regenerative medicine to generate desired cell types.

## Acknowledgements

We thank Nathaniel Roquet for helpful discussions.

## Funding

This work was supported by Korea Institute of Science and Technology (KIST) Institutional Programs (2E30240 to T.K.); Human Frontier Science Program (LT000595/2017-L to T.K.), and the Department of Defense (LC170525 W81XWH-18-1-0513 to T.K.L.)

## Author contributions

T.K., T.K.L. conceived the concept. T.K., T.K.L., B.W. and W.W. wrote the manuscript. T.K. performed all research. B.W. and W.W. provided the split recombinases used in the research. T.K.L. supervised the research.

## Data and materials availability

All data is available in the main text or the supplementary materials.

## Competing interests

T.K.L. is a co-founder of Senti Biosciences, Synlogic, Engine Biosciences, Tango Therapeutics, Corvium, BiomX, and Eligo Biosciences. T.K.L. also holds financial interests in nest.bio, Ampliphi, IndieBio, MedicusTek, Quark Biosciences, and Personal Genomics. Other authors declare no competing interests.

## List of Supplementary materials

**Materials and Methods**

**Figures S1-S3**

**Tables S1-S5**

**References (*29***)

## Notes

### Competing Interest Statement

The authors have declared no competing interest.

## References and Notes

1. M. Kondo, D. C. Scherer, A. G. King, M. G. Manz, I. L. Weissman, Lymphocyte development from hematopoietic stem cells. Current Opinion in Genetics & Development 11, 520–526 (2001).

2. C. A. Ortmann et al., Effect of Mutation Order on Myeloproliferative Neoplasms. New England Journal of Medicine 372, 601–612 (2015).

3. B. Vogelstein, K. W. Kinzler, The multistep nature of cancer. Trends in Genetics 9, 138–141 (1993).

4. N. Schönhuber et al., A next-generation dual-recombinase system for time- and host-specific targeting of pancreatic cancer. Nature Medicine 20, 1340–1347 (2014).

5. A. J. Meyer, T. H. Segall-Shapiro, E. Glassey, J. Zhang, C. A. Voigt, Escherichia coli “Marionette” strains with 12 highly optimized small-molecule sensors. Nature Chemical Biology 15, 196–204 (2019).

6. O. Borkowski, F. Ceroni, G.-B. Stan, T. Ellis, Overloaded and stressed: whole-cell considerations for bacterial synthetic biology. Current Opinion in Microbiology 33, 123–130 (2016).

7. M. Kafri, E. Metzl-Raz, G. Jona, N. Barkai, The Cost of Protein Production. Cell Reports 14, 22–31 (2016).

8. T. Frei et al., Characterization, modelling and mitigation of gene expression burden in mammalian cells. bioRxiv, 867549 (2019).

9. T. Miyamoto et al., Rapid and orthogonal logic gating with a gibberellin-induced dimerization system. Nat Chem Biol 8, 465–470 (2012).

10. F.-S. Liang, W. Q. Ho, G. R. Crabtree, Engineering the ABA Plant Stress Pathway for Regulation of Induced Proximity. Science Signaling 4, rs2 (2011).

11. B. H. Weinberg et al., High-performance chemical- and light-inducible recombinases in mammalian cells and mice. Nature Communications 10, 4845 (2019).

12. B. H. Weinberg et al., Large-scale design of robust genetic circuits with multiple inputs and outputs for mammalian cells. Nat Biotech 35, 453–462 (2017).

13. G. Meinke, A. Bohm, J. Hauber, M. T. Pisabarro, F. Buchholz, Cre Recombinase and Other Tyrosine Recombinases. Chemical Reviews 116, 12785–12820 (2016).

14. L. C. David et al., N availability modulates the role of NPF3.1, a gibberellin transporter, in GA-mediated phenotypes in Arabidopsis. Planta 244, 1315–1328 (2016).

15. A. G. Hinnebusch, I. P. Ivanov, N. Sonenberg, Translational control by 5′-untranslated regions of eukaryotic mRNAs. Science 352, 1413 (2016).

16. D. M. Shcherbakova, V. V. Verkhusha, Near-infrared fluorescent proteins for multicolor in vivo imaging. Nature Methods 10, 751 (2013).

17. X. Li et al., Generation of Destabilized Green Fluorescent Protein as a Transcription Reporter. Journal of Biological Chemistry 273, 34970–34975 (1998).

18. M. A. Li et al., Mobilization of giant piggyBac transposons in the mouse genome. Nucleic Acids Research 39, e148–e148 (2011).

19. T. B. Dolberg et al., Computation-guided optimization of split protein systems. bioRxiv, 863530 (2019).

20. Y. Gao, Y. Zhao, Self-processing of ribozyme-flanked RNAs into guide RNAs in vitro and in vivo for CRISPR-mediated genome editing. Journal of Integrative Plant Biology 56, 343–349 (2013).

21. K. Xie, B. Minkenberg, Y. Yang, Boosting CRISPR/Cas9 multiplex editing capability with the endogenous tRNA-processing system. Proceedings of the National Academy of Sciences 112, 3570 (2015).

22. D. J. H. F. Knapp et al., Decoupling tRNA promoter and processing activities enables specific Pol-II Cas9 guide RNA expression. bioRxiv, (2018).

23. L. Nissim, Samuel D. Perli, A. Fridkin, P. Perez-Pinera, Timothy K. Lu, Multiplexed and Programmable Regulation of Gene Networks with an Integrated RNA and CRISPR/Cas Toolkit in Human Cells. Molecular Cell 54, 698–710 (2014).

24. A. Chavez et al., Highly efficient Cas9-mediated transcriptional programming. Nat Meth 12, 326–328 (2015).

25. R. E. Haurwitz, S. H. Sternberg, J. A. Doudna, Csy4 relies on an unusual catalytic dyad to position and cleave CRISPR RNA. The EMBO Journal 31, 2824–2832 (2012).

26. J. E. Wilusz et al., A triple helix stabilizes the 3′ ends of long noncoding RNAs that lack poly(A) tails. Genes & Development 26, 2392–2407 (2012).

27. A. Hendel et al., Chemically modified guide RNAs enhance CRISPR-Cas genome editing in human primary cells. Nature Biotechnology 33, 985–989 (2015).

28. J. Drost et al., Sequential cancer mutations in cultured human intestinal stem cells. Nature 521, 43–47 (2015).

29. K. Clement et al., CRISPResso2 provides accurate and rapid genome editing sequence analysis. Nature Biotechnology 37, 224–226 (2019).

