## Supplementary material for "Scalable recombinase-based gene expression cascades": Methods, Supplemental figures and tables

### Materials and Methods

**Cell Culture.** HEK 293T cells were obtained from American Type Cell Culture, and were cultured in Dulbecco's Modified Eagle Medium (Gibco) supplemented with 10% fetal bovine serum (Corning), and penicillin/streptomycin (Gibco).

**GPC excision test.** HEK 293T cells were seeded in 96 well plates, and transfected 12-16 hours after seeding with unit GPC plasmid using Fugene HD (Promega). Ligands for CID are treated starting six hours after transfection for 24 hours. Cells were trypsinized for flow cytometry analysis using LSR Fortessa (BD Biosciences). The efficiency of excision was calculated as (% BFP positive)/ (%BFP or GFP positive) to account for transfection efficiency.

**Flow Cytometry.** Data obtained with LSR Fortessa (BD Biosciences) were compensated for spectral overlap and analyzed with FlowJo (FlowJo LLC.).

**Ligand decay assay.** HEK 293T cells were seeded in 96 well plates, and pretreated with CID ligands ($GA_3$-AM[Toronto Research Chemicals], $GA_3$ [Sigma], $GA_4$ [Sigma], ABA [Gold Biotechnology]). Twenty-four hours later, cells were washed with phosphate buffered saline and trypsinized for seeding in new 96 well plates in the absence of ligands. For zero hour wash, cells were seeded with DNA-Fugene HD mixture for reverse transfection. For other washout time points, cells washed out of ligands were trypsinized and seeded 12 hours before transfection. GIB-Cre and ABA-FlpO GPCs are transfected for GIB and ABA decay assays, respectively.

**DNA constructs.** Unit GPCs with optimized split recombinases are digested with PacI restriction enzyme (New England Biolabs), and was used to perform Gibson assembly with PCR amplicon containing the payload gene, 3x bovine growth hormone polyadenylation sequence, and 2x chicken hypersensitivity site 4 core sequence. The resulting unit GPCs were digested with SapI restriction enzyme (New England Biolabs) for golden gate assembly with SapI restriction enzyme and T4 DNA ligase (New England Biolabs). The complete list of plasmids used in this study is in Supplementary table 1. The protospacer sequences for sgRNAs are listed in Supplementary table 2.

**Reverse Transcription and quantitative PCR.** Total RNA from the cells were isolated with Trizol (Life Technologies) according to the manusfacturer's instructions. The resulting RNAs were reverse transcribed with MMLV reverse transcriptase (Promega). Complementary DNA were used as templates for quantification of genes using TOPreal qPCR 2x Premix (Enzynomics), and Applied Biosystems 7500 real-time PCR system. Primers used for quantitative PCR are listed in Supplementary table 3.

**T7 endonuclease assay.** Genomic DNA of HEK 293T cells were isolated using DNeasy Blood & Tissue Kit (Qiagen). Genomic loci flanking the sgRNA target sites were PCR amplified with Q5 DNA polymerase (New England Biolabs). The PCR product was purified with gel DNA recovery kit (Zymo Research Corp.). The purified PCR product was denatured at 95C for 5 minutes, and annealed by slowly cooling with ramp speed of -0.1C/s to 25C. The annealed PCR product was digested with 5U T7 endonuclease I (New England Biolabs) for 30 minutes in 37C for gel electrophoresis. Primers used for amplifying sgRNA target genomic loci are listed in Supplementary table 4.

**Next Generation Sequencing.** The PCR amplification product of size 200~300 base pairs flanking the sgRNA target sites were purified with gel DNA recovery kit (Zymo Research Corp.). The DNA library was prepared with Illumina TruSeq Nano DNA library Construction (insert size 350bp). The resulting DNA library was sequenced with HiSeq4000 (Illumina, 150nt paired-end). The next generation sequencing data was analyzed for indel frequency using

CRISPRESSO2 *(29)*. The primers used for generating amplicon for NGS analysis are listed in Supplementary table 5.

**Supplementary Text S1. Testing optimal CID ligand treatment schedule.**

We tested how quickly CID ligands can induce excision in a GPC. GIB treatment induced nearly complete excision of the GPC within 24 hours, with little additional excision achieved with longer duration of GIB treatment (Fig. S2A). To allow for sufficient expression of the next gene and degradation of the previous gene in the cascade, we treated cells with CID ligand for 48 hours for each stage in cascade.

We also examined the rate at which ligands were cleared from the cells to determine the minimum time course required for switching between ligands. The GIB molecule most commonly used for CID in mammalian cells is acetoxymethyl group modified gibberellin $A_3$ ($GA_3$-AM)(*9*), which can be trapped within cells by removal of the acetoxymethyl group by intracellular esterase. Other GIBs, such as $GA_4$, are known to be yeast membrane permeable without any modification (*14*) and may readily diffuse out of cells after ligand removal. As expected, $GA_3$-AM required more than 12 hours to be completely cleared from the cells while $GA_4$ was immediately cleared from the cells after ligand removal (Fig. S2B-C). Moreover, split-recombinase systems induced by $GA_4$ were as efficient as those induced by $GA_3$-AM (Fig. S2C). We subsequently used $GA_4$ to drive the gene expression cascade in the tandem GPC gene circuits throughout this study. We similarly tested the kinetics of ABA clearance from cells and found that cells were virtually free of ABA 12 hours after ligand removal (Fig. S2C). Thus, we utilized a ligand treatment schedule of 48 hours ligand treatment at each stage within the cascade, with a 12-hour gap between switching of the ligands.

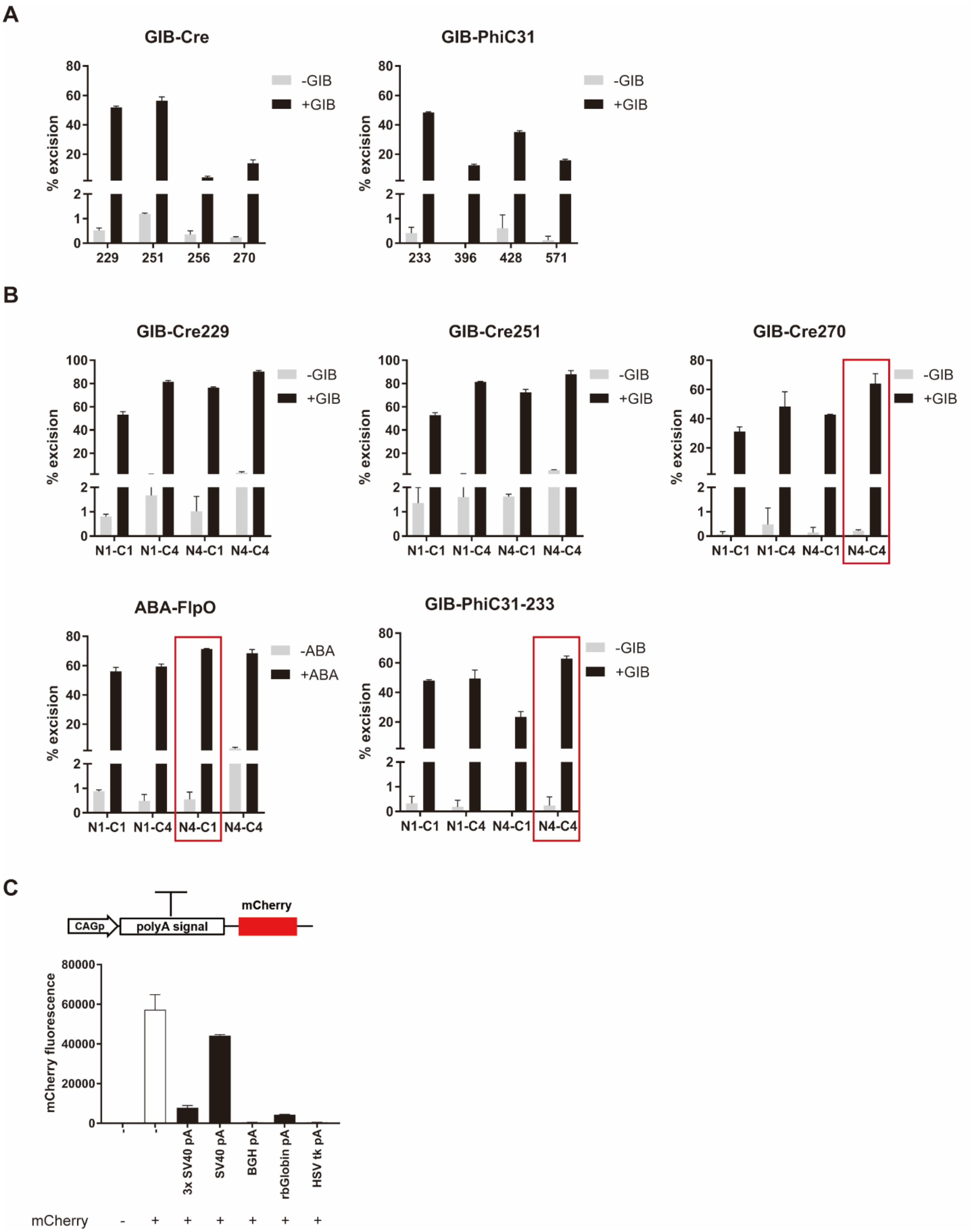


**Fig. S1. Optimization of unit GPCs.** (A-B) % BFP positive cells were calculated as % self-excision of GPCs with the experimental designs described in Figure 1B. (A) Determination of ideal recombinase split sites for minimal leakage in the absence of ligand and efficient ligand-induced self-excision of GPCs (n=2, mean± s.d.). (B) Determination of optimal number of nuclear localization signals (NLS) fused to each fragment of split recombinases for best activity in the presence of ligands (n=2, mean± s.d.). Nx-Cy indicates that the N-terminal fragment of recombinase was fused with x NLS(s), and the C terminal fragment was fused with y NLS(s). Red rectangles indicate the optimized split-recombinases used throughout this study. (C) Comparison of different polyadenylation signals for efficient transcription termination (n=2, mean± s.d.).

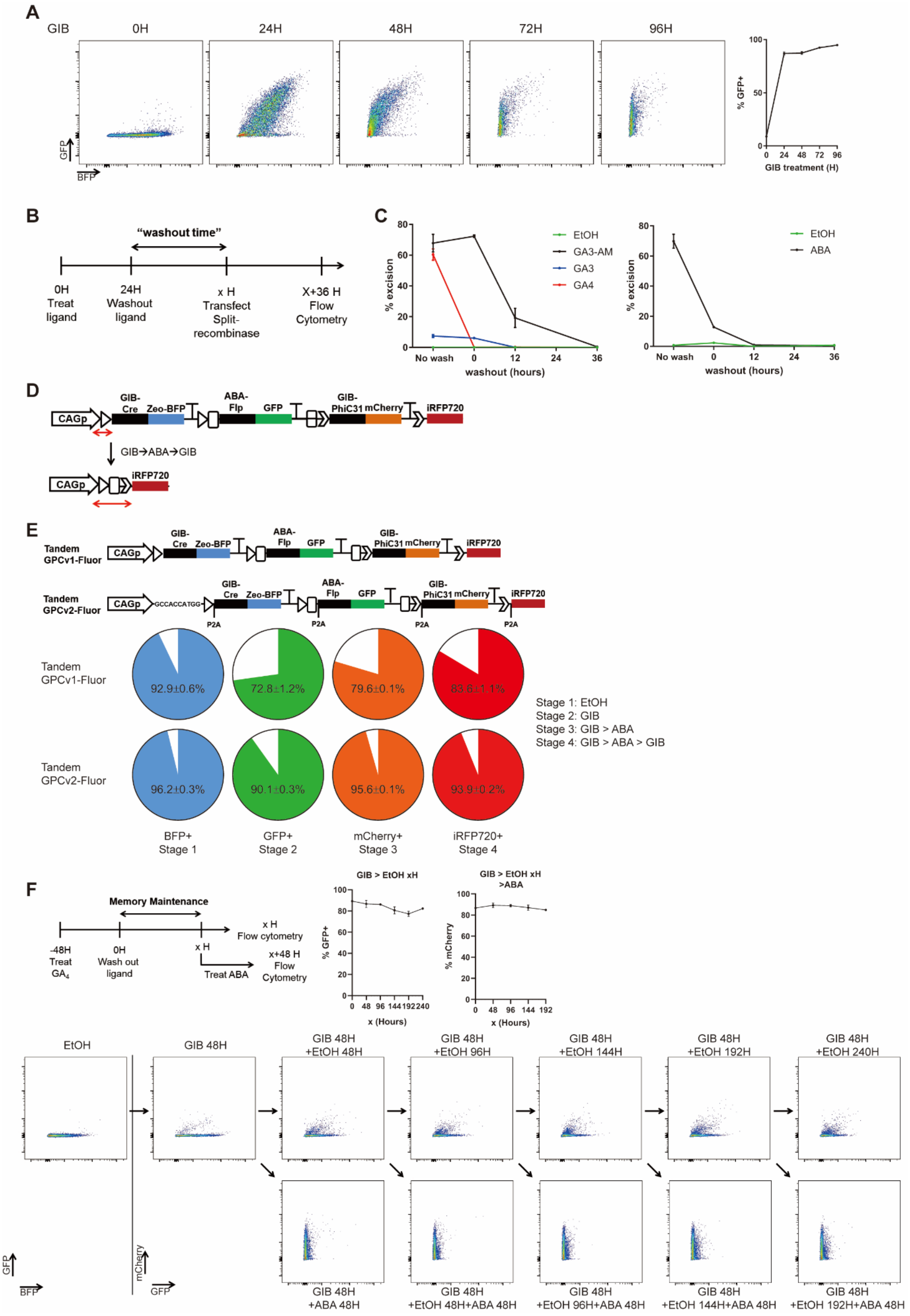

A
GIB
0H
24H
48H
72H
96H
GFP
BFP
% GFP+
GIB treatment (H)
B
"washout time"
0H Treat ligand
24H Washout ligand
x H Transfect Split-recombinase
X+36 H Flow Cytometry
C
% excision
EtOH
GA3-AM
GA3
GA4
ABA
No wash
washout (hours)
D
CAGp
GIB-Cre
Zeo-BFP
ABA-Flp
GFP
GIB-PhiC31
mCherry
iRFP720
GIB→ABA→GIB
E
Tandem GPCv1-Fluor
Tandem GPCv2-Fluor
GCCACCATGG
P2A
92.9±0.6%
72.8±1.2%
79.6±0.1%
83.8±1.1%
96.2±0.3%
90.1±0.3%
95.6±0.1%
93.9±0.2%
Stage 1: EtOH
Stage 2: GIB
Stage 3: GIB > ABA
Stage 4: GIB > ABA > GIB
BFP+ Stage 1
GFP+ Stage 2
mCherry+ Stage 3
iRFP720+ Stage 4
F
Memory Maintenance
-48H Treat GA4
0H Wash out ligand
x H
x H Flow cytometry
Treat ABA
x+48 H Flow Cytometry
GIB > EtOH xH
GIB > EtOH xH >ABA
% GFP+
% mCherry
x (Hours)
EtOH
GIB 48H
GIB 48H +EtOH 48H
GIB 48H +EtOH 96H
GIB 48H +EtOH 144H
GIB 48H +EtOH 192H
GIB 48H +EtOH 240H
GIB 48H +ABA 48H
GIB 48H +EtOH 48H+ABA 48H
GIB 48H +EtOH 96H+ABA 48H
GIB 48H +EtOH 144H+ABA 48H
GIB 48H +EtOH 192H+ABA 48H
mCherry
GFP

**Fig. S2. Optimization of tandem GPC circuit.** (A) Kinetics of recombinase-induced excision after ligand treatment. Tandem GPCv2-Fluor was treated with $GA_4$ for indicated durations. Excision was monitored as % GFP positive cells. (Left) Representative flow cytometry plots, (Right) Quantification of the % GFP positive cells (n=2, mean± s.d.). (B-C) Kinetics of CID ligand decay. (B) Cells were pretreated with various versions of gibberellin or abscisic acid. Gibberellin treated cells were transfected with plasmid encoding GIB-Cre GPC (used in figure 1D) at indicated time points after ligand washout. ABA treated cells were transfected with plasmid encoding ABA-Flp GPC (used in figure 1D) at indicated time points after ligand washout. (C) Quantification of the % excised cells with GPCs delivered after various washout time (n=2, mean± s.d.). (D) Lengthening of “scar” 5’-UTR sequences after each excision. The “scar” sequence is indicated with the red arrow. (E) Improved “tandem GPCv2” circuit design to prevent 5’-UTR lengthening. Pie chart below shows the fraction of cells expressing the desired payload gene after each indicated sequence of ligand treatment. (F) Memory maintenance by the gene expression cascade. (Top left) Experimental scheme. (Top right) Quantification of mean GFP level and fraction of GFP positive cells, and mean mCherry level after 48 hours of ABA treatment with fraction of mCherry positive cells (n=3, mean± s.d.) (Bottom) Representative flow cytometry plots.

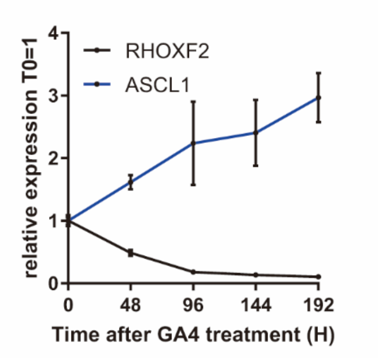


**Fig. S3. Optimization of tandem GPC circuit for sequential sgRNA expression.** Kinetics of sgRNA expression and decay after excision of GPC. Tandem GPCv2-CRISPRa was treated with gibberellin for the indicated durations. Relative expression level of the target genes RHOXF2, ASCL1, the target of first and second payload sgRNA, respectively, was quantified by RT-qPCR (n=2, mean± s.d.).

**Table S1.**
List of plasmids used in this study.

| Name | Description | Used in Figure |
|---|---|---|
| pTK1173 | EFS-LoxPGIBCre251-GFP-LoxP-BFP | S1A |
| pTK1174 | EFS-LoxPGIBCre229-GFP-LoxP-BFP | S1A |
| pTK1175 | EFS-LoxPGIBCre256-GFP-LoxP-BFP | S1A |
| pTK1176 | EFS-LoxPGIBCre270-GFP-LoxP-BFP | S1A |
| pTK1181 | EFS-LoxPGIBCre229-GFP-LoxP-BFP with 4x-1xNLS | S1B |
| pTK1182 | EFS-LoxPGIBCre229-GFP-LoxP-BFP with 1x-4xNLS | S1B |
| pTK1183 | EFS-LoxPGIBCre229-GFP-LoxP-BFP with 4x-4xNLS | S1B |
| pTK1184 | EFS-LoxPGIBCre251-GFP-LoxP-BFP with 4x-1xNLS | S1B |
| pTK1185 | EFS-LoxPGIBCre251-GFP-LoxP-BFP with 1x-4xNLS | S1B |
| pTK1186 | EFS-LoxPGIBCre251-GFP-LoxP-BFP with 4x-4xNLS | S1B |
| pTK1187 | EFS-LoxPGIBCre270-GFP-LoxP-BFP with 4x-1xNLS | S1B |
| pTK1188 | EFS-LoxPGIBCre270-GFP-LoxP-BFP with 1x-4xNLS | S1B |
| pTK1189 | EFS-LoxPGIBCre270-GFP-LoxP-BFP with 4x-4xNLS | S1B, S2C, 1C-E |
| pTK1224 | EFS-FRT ABAFlpO-GFP-FRT-BFP | S1B |
| pTK1225 | EFS-FRT ABAFlpO-GFP-FRT-BFP with 4x-1xNLS | S1B, S2C, 1C-E |
| pTK1226 | EFS-FRT ABAFlpO-GFP-FRT-BFP with 1x-4xNLS | S1B |
| pTK1227 | EFS-FRT ABAFlpO-GFP-FRT-BFP with 4x-4xNLS | S1B |
| pTK1196 | EFS AttP-GIBPhiC233-GFP-AttB-BFP | S1A |
| pTK1197 | EFS AttP-GIBPhiC396-GFP-AttB-BFP | S1A |
| pTK1198 | EFS AttP-GIBPhiC428-GFP-AttB-BFP | S1A |
| pTK1199 | EFS AttP-GIBPhiC571-GFP-AttB-BFP | S1A |
| pTK1204 | EFS AttP-GIBPhiC233-GFP-AttB-BFP with 4x-1xNLS | S1B |
| pTK1205 | EFS AttP-GIBPhiC233-GFP-AttB-BFP with 1x-4xNLS | S1B |
| pTK1206 | EFS AttP-GIBPhiC233-GFP-AttB-BFP with 4x-4xNLS | S1B, 1C-E |
| pTK1423 | EFS-LoxP-GIBCre270 Y324F GFP-LoxP-BFP with 4x-4xNLS | 1C-E |
| pTK1115 | PB CAG-3xSVpA-mCherry | S1C |
| pTK1116 | PB CAG-1xSVpA-mCherry | S1C |
| pTK1117 | PB CAG-BGHpA-mCherry | S1C |
| pTK1118 | PB CAG-rbGlobpA-mCherry | S1C |
| pTK1119 | PB CAG-HSVpA-mCherry | S1C |
| pTK1230 | PB CAG-2xSapI-iRFP w/o PEST for tGPC v1 | S2E |
| pTK1263 | PB CAG-iRFP720 w/o PEST for tGPC v2 | S2A, S2E-F, 2B-E |

| | | |
|---|---|---|
| pTK1442 | EFS-LoxPGIBCre270-ZeoBFP-3xBGH-2xcHS4 with 4x-4xNLS for tGPC v1 | S2E |
| pTK1443 | EFS-FRT-ABAFlpO-GFP-3xBGH-2xcHS4 with 1x-4xNLS for tGPC v1 | S2E |
| pTK1444 | EFS-AttP-GIBCphiC-mCherry-3xBGH-2xcHS4 with 4x-4xNLS for tGPC v1 | S2E |
| pTK1445 | EFS-LoxPGIBCre270-ZeoBFP-3xBGH-2xcHS4 with 4x-4xNLS for tGPC v2 | S2A, S2E-F, 2B-E |
| pTK1446 | EFS-FRT-ABAFlpO-GFP-3xBGH-2xcHS4 with 1x-4xNLS for tGPC v2 | S2A, S2E-F, 2B-E |
| pTK1447 | EFS-AttP-GIBCphiC-mCherry-3xBGH-2xcHS4 with 4x-4xNLS for tGPC v2 | S2A, S2E-F, 2B-E |
| pTK1312 | PB EF1a-dCas9-VPR-P2A-Csy4-CAG-28-sgTTN-28 | 3A |
| pTK1313 | PB EF1a-dCas9-VPR-P2A-CAG-tRNA$^{Gly\ C55G}$-sgTTN-tRNA$^{Gly}$ | 3A |
| pTK1314 | PB EF1a-dCas9-VPR-P2A-Csy4-CAG-20-sgTTN-20 | 3A |
| pTK1328 | PB EF1a-dCas9-VPR-P2A-CAG-HH-sgTTN-HDV | 3A |
| pTK1280 | FUW sgTTN-CMV$_{minP}$-RFP-CMV-Puro | 3A |
| pTK1448 | EFS-LoxPGIBCre270-Zeo-20-sgRHOXF2-20-3xBGHpA-2xcHS4 with 4x-4xNLS | S3, 3B-C |
| pTK1449 | EFS-FRT-ABAFlpO-20-sgASCL1-20-3xBGHpA-2xcHS4 with 1x-4xNLS | S3, 3B-C |
| pTK1450 | EFS-AttP-GIBCphiC-20-sgHBG-20-3xBGHpA-2xcHS4 with 4x-4xNLS | S3, 3B-C |
| pTK1451 | EFS-LoxPGIBCre270-Zeo-20-sgAPC-20-3xBGHpA-2xcHS4 with 4x-4xNLS | 3D-G, 4A-F |
| pTK1452 | EFS-FRT-ABAFlpO-20-sgMLH1-20-3xBGHpA-2xcHS4 with 1x-4xNLS | 3D-G, 4A-F |
| pTK1453 | EFS-AttP-GIBphiC-20-sgSMAD4-20-3xBGHpA-2xcHS4 with 4x-4xNLS | 3D-G |
| pTK1492 | EFS-GIBphiC-20-Puro-sgSMAD4-20-3xBGHpA-2xcHS4 with 4x-4xNLS w/o AttP | 4A-F |
| pTK1501 | PB CAG-Cas9-P2A-Csy4-P2A-2xSapI-sgTP53 | 3D-G |
| pTK1502 | PB CAG-dCas9-VPR-P2A-Csy4-P2A-2xSapI-sgTTN | S3, 3B-C |
| pTK1503 | PB AttP-CAG-Cas9-P2A-Csy4-P2A-2xSapI | 4A-F |
| pTK1601 | tGPC Fluor v1=pTK1230+1442+1443+1444 | S2E |
| pTK1602 | tGPC Fluor v2=pTK1263+1445+1446+1447 | S2A, S2E-F,2B-E |
| pTK1603 | tGPC-CRISPRa=pTK1502+1448+1449+1450 | S3,3B-C |
| pTK1604 | tGPC-CRISPR=pTK1501+1451+1452+1453 | 3D-G |
| pTK1605 | tGPC-AttP-CRISPR=pTK1503+1451+1452+1492 | 4A-F |

**Table S2.**

Protospacer sequences of sgRNA used in this study

| Gene | Sequence |
|---|---|
| RHOXF2 | AACGCGTGCTCTCCCTCATC |
| ASCL1 | CGGGAGAAAGGAACGGGAGG |
| HBG | CTTGACCAATAGCCTTGACA |
| TTN | CCTTGGTGAAGTCTCCTTTG |
| APC | TCTGTATAAATGGCTCATCG |
| MLH1 | TAATAGTAACATGAGCCACA |
| SMAD4 | TGTCCTTCAGTGGACAACGA |
| TP53 | CCATTGTTCAATATCGTCCG |

**Table S3.**
PCR primers for RT-qPCR.

| Gene | Sequence |
|---|---|
| RHOXF2 | F: GGCAAGAAGCATGAATGTGA |
| | R: GCCATTAATGCCCTCTGATG |
| ASCL1 | F: CGCGGCCAACAAGAAGATG |
| | R: CGACGAGTAGGATGAGACCG |
| HBG | F: AGATGCCACAAAGCACCTG |
| | R: CTGCAGTCACCATCTTCTGC |
| TTN | F: TGTTGCCACTGGTGCTAAAG |
| | R: ACAGCAGTCTTCTCCGCTTC |

**Table S4.**

PCR primers used for T7 endonuclease assay

| Gene | Sequence |
|---|---|
| APC | F: CCCTAGAACCAAATCCAGCA |
| | R: TATCATCCCCCGGTGTAAAA |
| MLH1 | F: GACCCCGTCATAGCACAGTT |
| | R: GCAAAGGGGCAGAAATTACA |
| SMAD4 | F: TTGGGTTGGGTTTCTAGAGG |
| | R: GTGGAAGCCACAGGAATGTT |
| TP53 | F: AGGGTGTGATGGGATGGATA |
| | R: CTGCCCTGGTAGGTTTTCTG |

**Table S5.**

PCR primers used for next generation sequencing analysis to quantify indel frequency. Six nucleotides barcodes "NNNNNN" are used for multiplexing.

| Gene | Sequence |
|---|---|
| APC | F: NNNNNNTCCAGGTTCTTCCAGATGCT |
| | R: TTTTTCTGCCTCTTTCTCTTGG |
| MLH1 | F: NNNNNNCCCTTTGGTGAGGTGACAGT |
| | R: CGTACTCAAGATCTCTGCCAAA |
| SMAD4 | F: NNNNNNTCAAGTATGATGGTGAAGGATGA |
| | R: CTTGTGGAAGCCACAGGAAT |
| TP53 | F: NNNNNNCTGCCCTGGTAGGTTTTCTG |
| | R: CCTGGTCCTCTGACTGCTCT |
